# Proteome of human glioblastoma and meningioma tissue small extracellular vesicles

**DOI:** 10.1101/2024.04.15.589661

**Authors:** Huaqi Su, Adityas Purnianto, Andrew H. Kaye, Andrew P. Morokoff, Katharine J. Drummond, Stanley Stylli, Laura J. Vella

## Abstract

Small extracellular vesicles have gained attention in neuroscience due to their role in cell-to-cell communication and their potential diagnostic and therapeutic applications. Despite progress in the field, there remains a gap in our understanding of the composition and function of extracellular vesicles with regards to brain tumours. Previous studies have primarily evaluated extracellular vesicles obtained from patient fluids or cell culture medium, rather than directly from tumour tissue.

Here we successfully isolated small extracellular vesicles from surgical tissue biopsies of glioblastomas or meningiomas, marking, marking the first report of *in situ* extracellular vesicle isolation from brain tumours. The protein content of the tumour tissue and their extracellular vesicles was characterized using tandem mass spectrometric proteomics, revealing proteins exclusively detected or enriched in extracellular vesicles relative to the tumour tissue. While our study confirmed proteins previously identified in glioblastoma and meningioma extracellular vesicles from various sources, it also identified novel proteins and pathways associated with extracellular vesicles from these tumour types.

This study underscores the benefit of analysing *in situ* extracellular vesicles derived directly from brain tissue for insights into tumour biology and highlights the need for further research comparing extracellular vesicles from various types and grades of brain tumours.

## Introduction

Small extracellular vesicles (EVs) have gained attention in neuroscience owing to their involvement in intercellular communication and potential applications in both diagnosis and treatment of disease (1, 2). These small membrane-bound vesicles, released by both cancer and non-cancerous cells, carry molecules between cells, including proteins, lipids, and nucleic acids and have demonstrated roles in neural development and activity. The intrinsic role of EVs in the neural network means they have become a focus of studies on^i^ brain pathologies including glioblastoma (GBM), where they are proposed to have multifaceted roles in tumour progression, metastasis, immune modulation, and the development of drug resistance (3, 4)

Due to the presence of EVs in the extracellular environment they are accessible for non-invasive studies, and thus there is intense interest in using EVs isolated from bodily fluids (including blood, cerebrospinal fluid (CSF) and urine)to investigate tissue and disease specific biomarkers. By analysing the content of EVs, researchers have been able to identify specific biomarkers associated with different tumour types, as well as monitor disease progression and treatment response (5, 6). Defining the cargo in tumour tissue EVs offers a window into the molecular underpinnings of disease. Understanding this cargo can reveal insights into tumour biology and discover novel therapeutic targets.

Despite progress in understanding the significance of EVs in brain function and the genesis and progression of brain tumours, there is a gap in our knowledge. The majority of prior investigations have primarily been on EVs obtained from patient fluids or cell cultures (4, 7–12), rather than directly from the tumour tissue itself, which is important as these EVs may not adequately represent the complexity and specificity of those in tumour tissue. Relative to the routine isolation of EVs from fluids, including conditioned cell culture media, blood, urine or CSF, many technical issues must be addressed when isolating EVs directly from solid tissue. Overcoming the technical challenges in isolating EVs directly from tumour tissue may provide more accurate and direct insights into tumour biology.

Recent advancements, including methods for isolating and enriching EVs from human brain tissue, represent a significant step forward (13, 14). Using these methods, which we have previously developed for the isolation and enrichment of EVs from human brain tissue (13, 14), here we have, for the first time, isolated EVs from human meningioma (MNG), a tumour arising from the from the meninges covering the brain and spinal cord and glioblastoma (GBM), a highly aggressive glial brain tumour. We analysed the composition of these EVs using mass spectrometry-based proteomics and offer insights into MNG and GBM EV function and potential therapeutic targets.

## Materials and Methods

### Human tumour tissue

GBM and MNG tissue biopsies (fresh frozen at -80°C) were obtained from the Royal Melbourne Hospital (RMH) Neurosurgery Brain and Spine Tumour Tissue Bank (Human Research Ethics Committee Approval: HREC 2001.085) and examined according to the approved protocols in project (HREC2019.082). Individual patient demographic and clinical information was collected (**Table 1)**.

### EV enrichment from tumour tissue

The EV isolation method was modified from our previously published protocol (13–15). A schematic of the workflow is shown in **Supplemental Figure 1**. Frozen GBM (n = 8) and MNG (n = 8) tissues were sliced lengthways on ice to generate 1–2 cm long, 2–3 mm wide tissue sections. Tumour tissue pieces of approximately 30 mg from each patient biopsy (“Brain Total”) were collected, weighed and placed in a 19x volume by tissue weight of Dulbecco’s phosphate buffered saline solution (DPBS, Thermo Fisher Scientific) containing 1x PhosSTOP^TM^ phosphatase inhibitor (Sigma Aldrich) / cOmplete^TM^ protease inhibitor (including EDTA, Sigma Aldrich) for immunoblot analysis. The remaining cut tissue sections were weighed and incubated with 50 U/mL collagenase type 3 digestion buffer (#CLS-3, CAT#LS004180, Worthington) at ratio of 8 μL /mg tissue in a shaking water bath (25°C, 20 mins). After 15 mins, the solution was gently pipetted up and down once and then incubated for a further 5 mins, which was followed by the addition of ice-cold 10x inhibition buffer (10x phosphatase inhibitor and 10x protease inhibitor in DPBS) resulting in a 1X working solution.

The dissociated tissue in solution was subjected to a series of centrifugations, including 300 x g, 4°C for 5 min, 2000 x g, 4°C for 10 min and 10,000 x g, 4°C for 30 min. The 10,000 x g spin supernatant was loaded onto the qEV10 70 nm size exclusion column (SEC) column (IZON Science), and EV elution (20 mL) were collected after the void volume (20 mL) following the manufacturer’s instruction. The 20 mL EV solution was subjected to ultracentrifugation for 40 min using a 10 kDa ultra centrifugal filter (UFC8010, Millipore). Aliquots of 5 μL and 50 μL were sampled for transmission electron microscopy (TEM) and nanoparticle tracking analysis (NTA), respectively. In addition, a 100 μL EV aliquot was collected for protein quantitation and proteomic analysis.

### Tissue homogenization and protein quantification

Tumour tissue in DPBS solution with 1x phosphatase and protease inhibitor was homogenised for 12 sec at 50% intensity using a tapered microtip attached to the Sonifier Cell Disruptor (Branson) and sonicated in an ice-cold water bath for 20 min. A 100 μL aliquot of tumour homogenate from each patient sample was centrifuged at 10,000 x g, 4°C for 5 min and the supernatant was mixed with a 20% SDS solution to achieve a protein solution in 5% SDS, prior to heating at 95°C for 10 min. The 100 μL concentrated EV solutions were sonicated in an ice-cold water bath sonicator for 20 min. The protein concentration in the tissue homogenates and EV suspensions was determined with a Pierce^TM^ BCA assay kit (Thermo Fisher Scientific) according to the manufacturer’s instruction.

### SDS-PAGE and Western Blot (WB) analysis

Samples were prepared in 4 x Laemmli sample buffer, then boiled (10 mins, 90°C) followed by centrifugation (14,000 x g, 1 min). Normalised samples (1 – 3 µg) were electrophoresed on the 4-20% Criterion TGX Stain-Free Precast gels (BioRad) in Tris/Glycine/SDS running buffer (BioRad) for 30 mins at 245 V. The proteins were transferred onto nitrocellulose membranes using an iBlot^TM^ 2 Dry Blotting System (P0 method, Thermo Fisher Scientific). Susbequently, the membranes were blocked with 5% (w/v) skim milk in TBS-T (1 hr, room temperature) followed by an overnight incubation with the following primary antibodies - (hexokinase-1 #2024 from Cell Signalling Technology, flotillin-1 # #610821 from BD Biosciences, syntenin #ab133267 from Abcam, TSG101 #T5701 from Sigma) at 4°C. Membranes were subsequently washed four times with TBS-T (30 min, RT) following a 1 hr, RT incubation with the secondary anti-rabbit antibodies, the IRDye® 800CW Goat anti-Rabbit IgG secondary antibody (#925-32211, LICOR) or the IRDye® 680CW Goat anti-Mouse IgG secondary antibody (#926-68070, LICOR). All antibodies were diluted in 5% (w/v) skim milk in TBS-T. Membranes were washed four times with TBS-T (30 min, RT). The membranes were imaged on an Odyssey® Fc Imaging System (LI-COR).

### Transmission electron microscopy (TEM)

A 5 μL EV suspension fixed in 1% (w/v) electron microscopy-grade glutaraldehyde was absorbed onto neutralised 300-mesh carbon-coated formvar copper grids (ProSciTech) for 1 min. Excessive liquid was removed and grids were washed twice in µL of dH_2_O, and negatively stained by incubating twice in 2% (w/v) saturated aqueous uranyl acetate for 1 min. Excessive stain was removed, and grids were air dried. Images were taken on a FEI Tecnai F20 (FEI, Eindhoven) transmission electron microscope. Wide field images encompassing multiple vesicles were captured to provide an overview of the fraction in addition to close-up images. Electron microscopy was performed in the Ian Holmes Imaging Centre at The University of Melbourne.

### Nanoparticle tracking analysis (NTA)

50 μL EV suspensions (50ml) were diluted with 0.22 µM filtered MilliQ water to obtain a particle concentration of 100-200 particles per frame, as determined by the NanoSight NS300 instruments (Malvern Instruments). For each sample, five videos of 30 seconds were acquired for each measurement using a syringe pump speed of 40, while the detection threshold was set at 4 and camera level was 12. Experimental videos were analysed by NTA 3.2 Dev Build 3.2.16 software (Malvern). NTA was performed in the Materials Characterisation and Fabrication Platform at The University of Melbourne.

### Proteomics sample preparation

20 µg of protein from “Brain Total” and EV solution in 5% SDS were transferred to protein low bind tubes, reduced and alkylated by 10 mM tris (2-carboxymethyl) phosphine (TCEP) and 40 mM chloroacetamide (CAA) in 50mM triethylammonium bicarbonate (TEAB) buffer at 99°C for 5 min and the reaction was then quenched in 1.2% phosphoric acid. A 7X volume of binding buffer (0.1 M TEAB and 90% methanol solution, pH 7.1) was added to each sample to generate the protein particles. The protein suspensions were transferred to micro S-Trap cartridges (Protifi) followed by centrifugation at 4,000 x g for 1 min to remove solvent and trap protein particles. Protein bound to the quartz membrane was washed with 150 µL binding buffer three times and digested with Pierce™ Trypsin Protease MS-Grade (Thermo Scientific) dissolved in 50 mM TEAB buffer with a digestion ratio of 1:20, trypsin:protein, w:w. Digestion of the tissue was performed overnight in a 37°C incubator. Digested peptides were eluted with three subsequent buffers, 40 µL of 50 mM TEAB buffer, 40 µL of 0.2% formic acid and 40 µL of 50% aqueous acetonitrile (ACN) containing 0.2% formic acid. Protein solution was dried and reconstituted in 80 µL of 2% ACN containing 0.05% trifluoroacetic acid.

### Data dependent analysis (DDA) tandem mass spectrometric (MS/MS) proteomics

Samples were processed using the nano-LC system and the Ultimate 3000 RSLC (Thermo Fisher Scientific) maintained at 50°C. A total of 1 µg of peptides was loaded onto the Acclaim PepMap nano-trap column (C18, 100 Å, 75 μm × 2 cm, Thermo Fisher Scientific) with an isocratic flow of 6 µL/min of 2% ACN containing 0.05% TFA for 6 min. Then, the nano-trap column was switched to the Acclaim PepMap RSLC analytical column (C18, 100 Å, 75 μm × 50 cm, Thermo Fisher Scientific). Solvent A was 0.1% v/v formic acid and 5% v/v dimethyl sulfoxide (DMSO) water solution, and solvent B was ACN with 0.1% v/v formic acid and 5% DMSO water solution. The peptides were eluted with a flow rate of 0.3 μL/min. The gradient increased from 2% B to 23% B for 89 min, 23% B to 40% B for 10 min, 40% B to 80% B for 5 min, maintained at 80% B for 5 min, then dropped to 2% immediately and stabilized at 2% for 10 min until the experiment finished.

The MS experiments were performed on an Orbitrap QExactive Plus mass spectrometer (Thermo Fisher Scientific) using a nano-ESI ion source in positive ionisation mode with a spray voltage of 1900V. The ion transfer tube was heated to 250°C. MS and MS/MS data were acquired with full scan MS spectra acquired with a 3 second cycle time at the range of 375-1400 m/z, orbitrap resolution of 120,000, an automatic gain control (AGC) value of 400,000, a maximum injection time of 50 ms and an RF lens at 30%. Settings were data-dependent of 5e4, isolation window of 1.6 m/z, collision energy of 30%, resolution of 15,000, an AGC value of 50,000 and a maximum injection time of 22 ms and 30s dynamic exclusion time. Experiments were performed at the mass spectrometry and proteomics facility (MSPF) located in the Bio21 Institute, Parkville, VIC, Australia.

### Protein identification

Peptide and protein identification was performed with the MaxQuant platform (version 1.6.17.0), searching against the FASTA sequences of the Swiss-Prot Homo Sapiens proteome database (download from uniport in June 2021, containing 20,380 reviewed proteins). In the label free quantification (LFQ) analysis, the identification of peptides was conducted with the minimum peptide length of 7, maximum peptide mass of 4000 Da. Variable modifications of Oxidation (M) and a fixed modification of carbamidomethylation of cysteine were applied. Precursor ion tolerance was set at 20 ppm. Digestion was trypsin/P with a max missed cleavage of 2. A false discovery rate (FDR) of 0.01 was applied at both the peptide and protein levels. The identification of protein required razor + unique peptides number larger or equal to one and intensities of razor + unique peptides were used for protein quantification. The intensity based absolute quantitation (iBAQ) was checked for quantitation. Others followed default settings.

### Data analysis and statistical analysis

Proteomic data were further analysed using the Perseus software (version 1.6.15.0). Proteins present at above 70% samples in at least one group were included for downstream comparison. Potential contaminants were removed. Comparison of the LFQ values of the selected protein markers was performed and tested with student *t* test in GraphPad Prism 9. Comparison of tumours and the derived EV was performed and tested with Welch’s T-test with *post-hoc* Benjamini-Hochberg FDR multiple comparison correction (FDR was set at 0.01) in Perseus. Volcano plots were generated using GraphPad Prism 9. After searching and filtering, enrichment analyses for gene ontology, KEGG pathways and Reactome pathway for small EVs enriched fractions were performed using GeneCodis4 with default significance threshold. String Network analysis was performed with high confidence 0.7 using K means clustering.

The spatial expression profile of the GBM EV exclusive and enriched proteins was examined using the Ivy GAP Glioblastoma Atlas (16) and their impact on GBM patient survival determined using the Glioma Bio Discovery Portal (17).

## Results

### Characterization of EVs isolated from human brain tumours

We previously reported a method to isolate small EVs from the extracellular matrix (ECM) of post-mortem human frontal cortex (13, 15, 18). The vesicles contain the hallmarks of endosomal derived exosomes (13, 15) and fulfilled the experimental requirements of the International Society for Extracellular Vesicles 2018 guidelines (19). Dissociated collagenase treated GBM and MNG tissue (brain total) was subject to sequential centrifugation and SEC to isolate EVs from the ECM of the tumour tissue. All EVs were subject to WB and TEM (**Figure 1**) to screen for EV markers and contaminants before subjecting samples to downstream proteomic analysis. Relative to the GBM and MNG tumour tissue homogenate, EV samples were enriched in the small EV marker syntenin and deplete of hexokinase (**Figure 1A**). EVs contained flotillin, while TSG101 was not detected. TEM images showed small, cup-shaped membrane vesicles with a diameter of 40-200 nm for both GBM and MNG samples (**Figure 1B and 1C**). The TEM was supported by NTA which demonstrated that GBM EVs had a mean diameter of 158.6 nm ± 64.8 nm, and MNG EVs of 160.5 nm ± 64.6 nm (**Figure 1D and 1E**). Together, these results suggest that the vesicles isolated from human GBM and MNG tissue are consistent with the morphology, size and protein co-enrichment of small EVs, as we have previously reported (13, 15).

**Figure 1.**
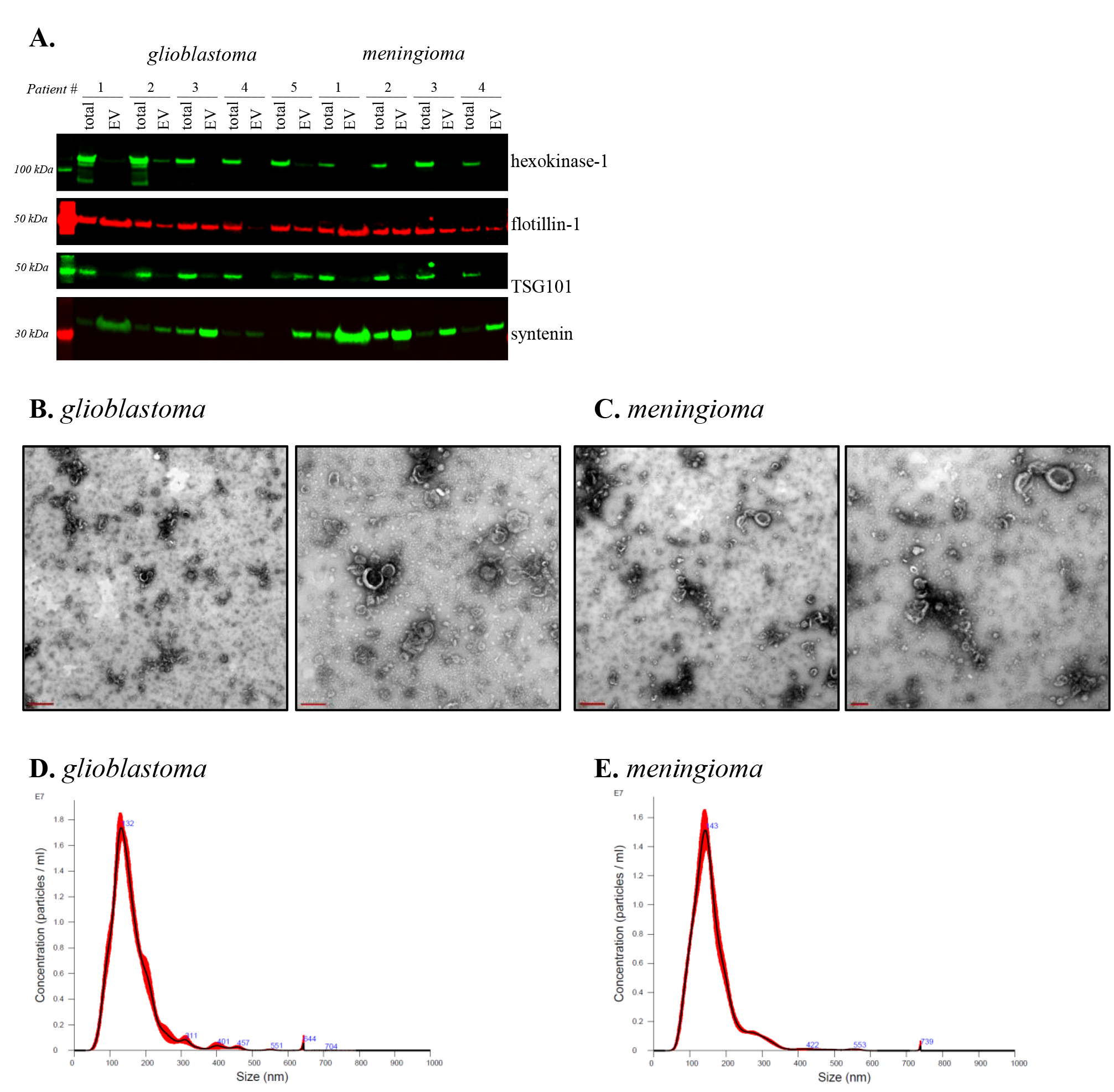
Characterization of EVs isolated from human glioblastoma and meningioma tumour tissue. (**A**) Western blot analysis. Equivalent amount of protein (6.5 μg) from human tumour homogenate (brain total), or TDEV were subjected to SDS-PAGE. Stain-free technology was used to confirm equal loading. Tumour homogenates, were enriched in hexokinase-1, while this protein was either not detectable or present in low amounts in an equivalent amount of EV protein. Proteins typical of endosome derived exosomes, TSG101 and syntenin, were observed in BDEVs, illustrating that EVs may be enriched in exosome- like vesicles. Immunoblots images are representative of the 5 GBM and 4 MG patient samples. Transmission electron microscopy (TEM) of the (**B**) glioblastoma EVs and **(C)** meningioma EVs provides an overview with scale bar representing 500 nm (left) and the close-up image (right) shows clearer small, cup-shaped EVs which is consistent with the morphology of exosomes with scale bar representing 200 nm. Nanoparticle tracking analysis (NTA) of the (**D**) glioblastoma EVs with a mean diameter of 158.6 nm ± 64.8 nm, and **(E)** meningioma EVs with a mean diameter of 160.5 nm ± 64.6 nm (SD).

Proteomics was used to further determine the protein content of tumour tissue and EVs isolated from the tissue (**Figures 2 – 6, Supplementary Tables 2 - 15**). The classical EV markers, syntenin and CD63 were enriched in GBM EVs, while CD81, CD63 and flotillin were enriched in MNG EVs (**Figure 2**). Analysis of gene ontology (GO) terms (20) revealed EVs from both tumour types were highly enriched in proteins associated with the cellular component terms ‘cytoplasm’, ‘exosomes’ and ‘lysosome’ highlighting the origin and extracellular nature of the vesicles (**Figure 2**).

**Figure 2.**
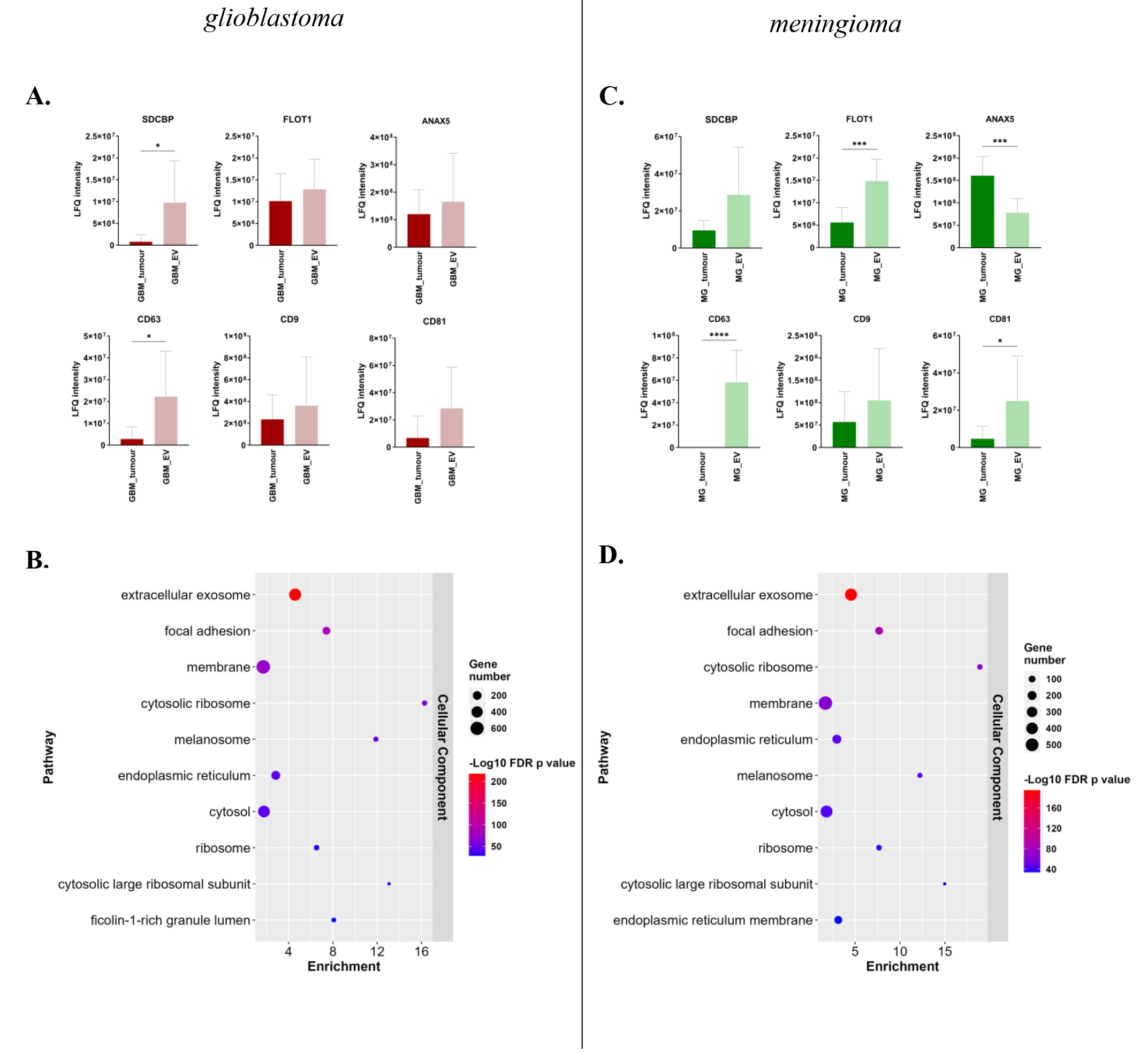
Proteins extracted from either tumour tissue or EVs isolated from said tumour tissue were submitted to proteomic analysis. (**A**) LFQ intensity graphs demonstrating enrichment of EV associated proteins in EVs isolated from GBM tumour tissue and (**B**) cellular component enrichment analysis of GBM EVs (882). (**C**) LFQ intensity graphs demonstrating enrichment of EV associated proteins in EVs isolated from MNG tumour tissue and (**D**) cellular component enrichment analysis of MNG EVs (812).

### Proteins exclusively associated with EVs relative to tumour

While the proteome of EVs comprises a subset of the parent tissue proteome, it is widely recognized that cells actively sort specific proteins into EVs to fulfill various functions. We compared the proteome of EVs with that of tumour tissue to determine which proteins were common between tumour tissue and EVs and which proteins were exclusively detected in EVs (**Figure 3**).

**Figure 3.**
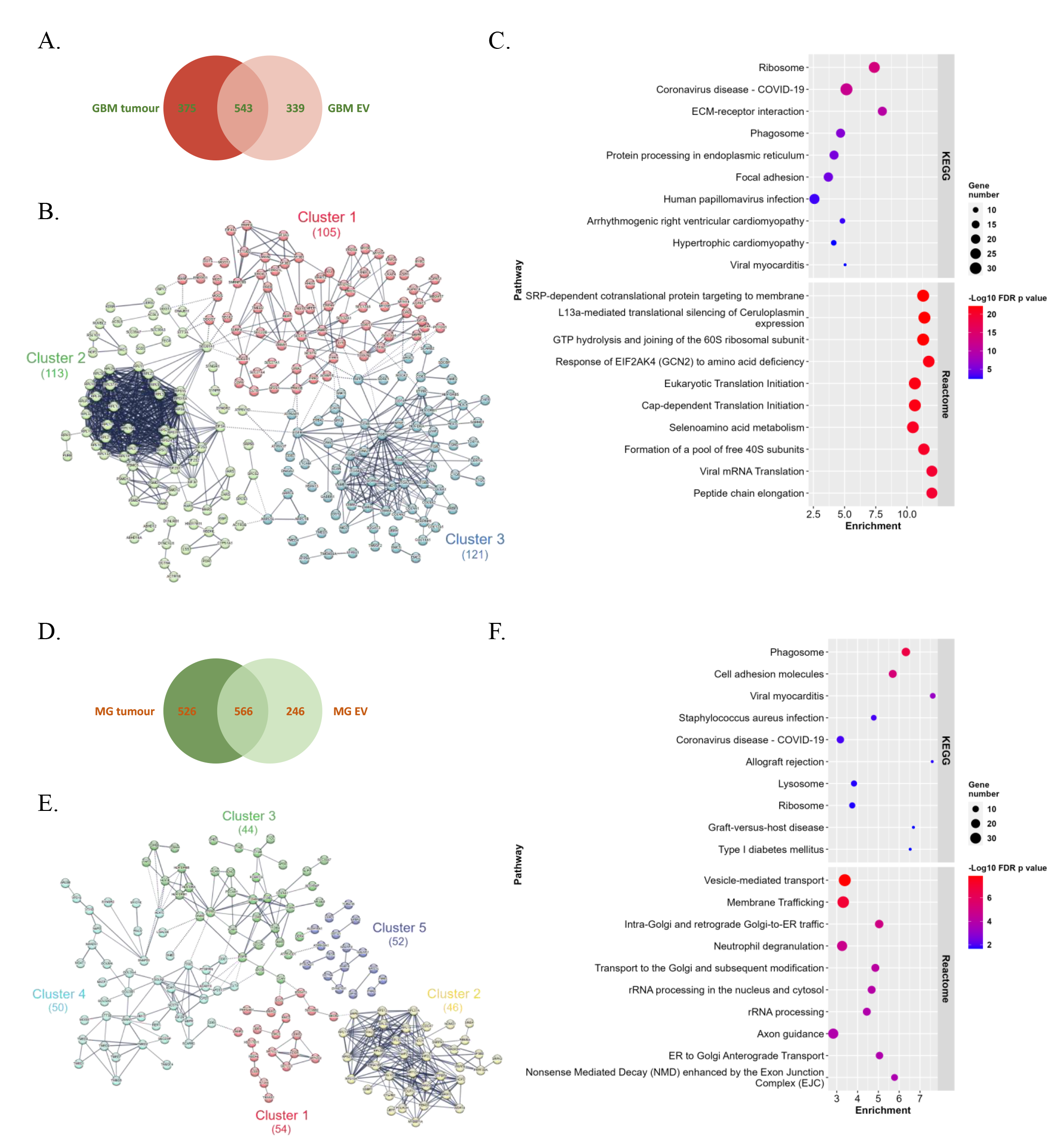
Analysis of proteins exclusive to EVs. (**A).** A two-way venn diagram showing proteins identified in GBM tumour and GBM EV with 339 proteins exclusive to GBM EV. (**B)**. String Network analysis of the 339 exclusive GBM EV proteins showing 3 clusters (high confidence 0.7, K means clustering used with 3 clusters, disconnected nodes hidden). Cluster 1: Proteosome/ribosome (105), cluster 2: ECM-receptor interaction (113), and cluster 3: synaptic vesicles (105). ©. Top 10 KEGG pathway enrichment and Top 10 Reactome enrichment of the 339 exclusive GBM EV proteins. **(D).** A two-way venn diagram showing proteins identified in MG tumour and MG EV with 246 proteins exclusive to MG EV. (**E)**. String Network analysis of the 246 exclusive MG EV proteins showing 5 clusters (high confidence 0.7, K means clustering used with 5 clusters, disconnected nodes hidden). Cluster 1: sphingolipid/sphinganine metabolism (54), cluster 2: ribosome (46), cluster 3: integrins (44), cluster 4: tetrespannin enriched/N-glycan biosynthesis (50), cluster 5: endoplasmic reticulum network/vesicle transport (52). (**F).** Top 10 KEGG pathway enrichment and Top 10 Reactome enrichment of the 246 exclusive MG EV proteins.

We detected 918 proteins in GBM tumours and 882 in the EVs isolated from these tissues (**Figure 3A and Supplementary Tables 2 - 4**). A common subset of 543 proteins were present in GBM tissue and EVs, while there were 339 proteins identified exclusively in GBM EVs. The proteins detected in GBM EVs (and not the corresponding tissue) include those involved in EV biogenesis such as CD63, SDCBP, VAMP2, CD44 (**Supplementary Table 5**). Other proteins identified exclusively in GBM EVs include proteins previously identified in GBM patient plasma derived EVs including Willebrand Factor (VWF) (21), the GBM surfaceome signature (HLA-DRA, CD44, EGFR and ITGB2) (22), CHI3L1 a serum biomarker of GBM (23), laminin subunit alpha-4 (LAMA4) which is increased in the CSF of GBM patients (24) and APP, ITGB1, IGFR2 which have been identified as EV markers of aggressive GBM (25) (**Supplementary Table 5**). We next analysed the protein interaction

network, conducted a Reactome analysis, and performed KEGG pathway analysis to determine the possible molecular mechanisms and signalling pathways that underlie the functions of EVs (GBM **Figure 3B-C**). Protein–protein interactions between the GBM EV exclusive proteins were investigated using the STRING database for functional protein association networks (https://string-db.org, version 11.0) (Szklarczyk et al., 2017). STRING analysis of exclusive GBM EV proteins identified three different clusters; 1: Proteosome/ribosome (105 proteins), 2: ECM-receptor interaction (113 proteins), and 3: synaptic vesicles (121 proteins) (**Figure 3B**). KEGG analysis identified terms such as ribosome, ECM-receptor interaction and focal adhesion but also terms associated with viral infection (**Figure 3C**). The Reactome pathway database identified pathways predominantly involved in translation (**Figure 3C**).

We detected 1092 proteins in the MNG tumour and 812 in MNG EVs (**Figure 3D and Supplementary Tables 6 - 8**). A common subset of 566 proteins were present in MNG tissue and EVs, while there were 256 proteins identified exclusively in MNG EVs (**Supplementary Table 9**). The proteins detected exclusively in MNG EVs include those involved in EV biogenesis (CD63, SDCBP, VAMP2, CD44) and PTGFRN which has previously been identified as a marker of meningioma EVs (8) (**Supplementary Table 9**). STRING analysis of exclusive MNG EV proteins identified five different clusters. Cluster 1: sphingolipid/sphinganine metabolism (54 proteins), cluster 2: ribosome (46 proteins), cluster 3: integrins (44 proteins), cluster 4: tetraspannin enriched/N-glycan biosynthesis (50 proteins) and cluster 5: endoplasmic reticulum (ER) network/vesicle transport (52 proteins) (**Figure 3E**). KEGG analysis identified terms such as phagosome, cell adhesion molecules, lysosome and ribosome (**Figure 3F**). The Reactome pathway database identified pathways predominantly involved in transport and trafficking including vesicle-mediated transport, membrane trafficking, golgi-to-ER trafficking (**Figure 3F**).

### Proteins enriched in EVs relative to tumour

We next performed differential expression analyses to determine the relative abundance of proteins common to tissue and EVs (**Figure 4 and 5 and Supplementary Tables 10 - 11**). We determined the differences in the abundance of the 543 proteins shared by GBM tumours and their EVs (**Figure 4A and Supplementary Table 10**). We identified 97 proteins that were significantly upregulated in GBM EVs (**Figure 4A**). These include MVP, ACTR3, PRKDC, APOE, TNC, MACF1, PDIA3 and RPS6 which have been previously identified in the cargo of GBM exosomes and are suggested drivers of invasion and or associated with more aggressive disease (25–32) (**Supplementary Table 10**). A protein-protein interaction enrichment analysis was carried out on the 339 exclusive and 97 significantly upregulated proteins in GBM EVs and the subnetworks analysed using the Molecular Complex Detection (MCODE) algorithm in Metascape (33) to identify densely connected network components. Five protein complex hubs were identified (MCODE 1 - 5). Proteins in MCODE 1 were significantly enriched in ribosome related terms including *SRP-dependent co-translational protein targeting to membrane, ribosome* and *cytoplasmic ribosomal proteins*. Proteins in MCODE 2 and MCODE4 were significantly enriched in endoplasmic reticulum terms including *protein processing in endoplasmic reticulum, response to endoplasmic reticulum stress* and *calnexin/calreticulin cycle,* and MCODE4 significantly enriched in *endoplasmic-reticulum-associated protein folding and degradation* terms and MCODE 5 enriched in RHO GTPase terms.

**Figure 4.**
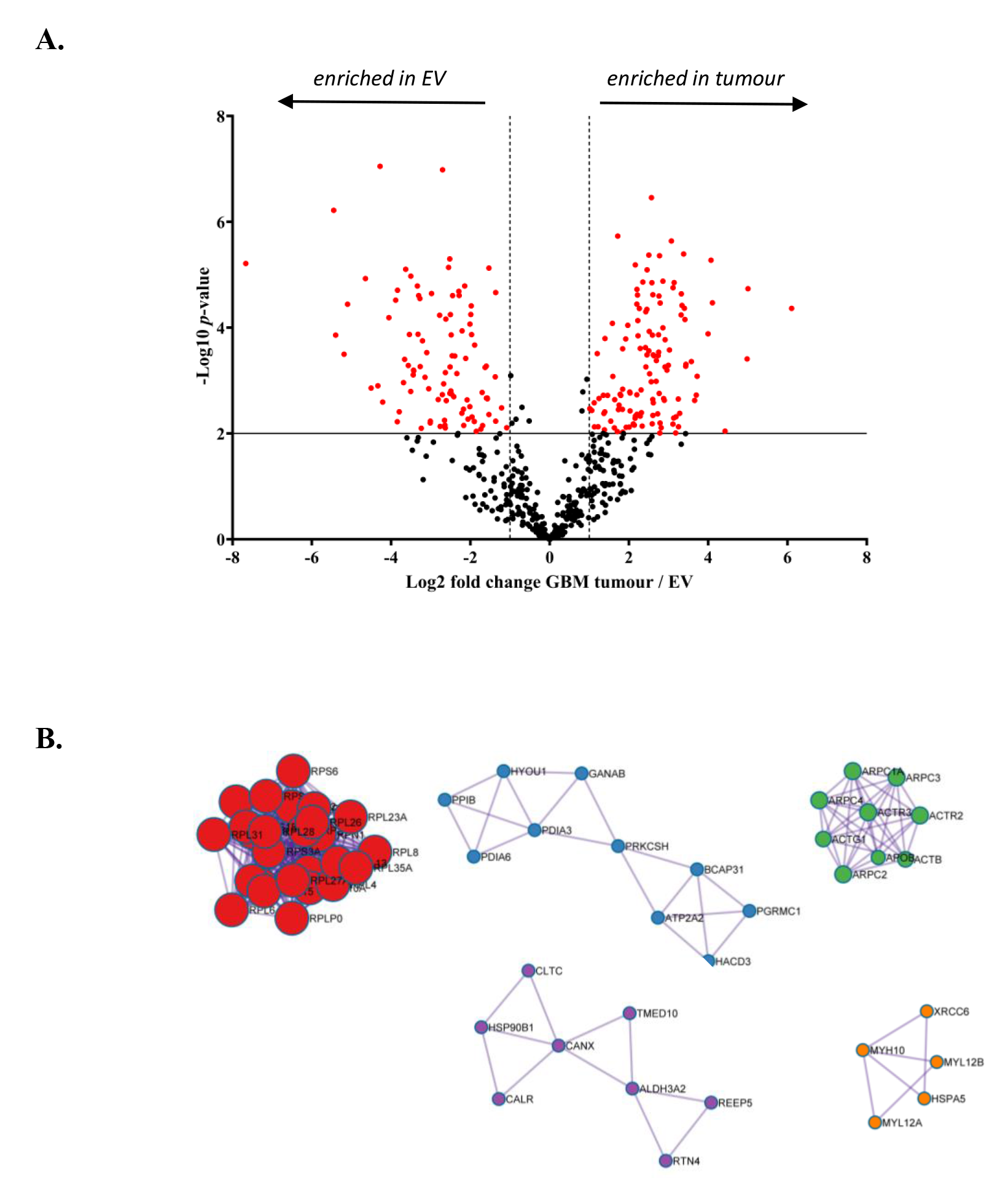
Analysis of proteins common between GBM tumours and their EVs. (**A).** Volcano Plot illustrating 97 proteins significantly enriched in GBM EVs. Comparison of GBM tumours and the derived EV was performed and tested with Welch’s T-test with post- hoc Benjamini-Hochberg FDR multiple comparison correction (FDR was set at 0.01). Volcano plot was generated using GraphPad Prism 9. **(B).** MCODE analysis of exclusive and significantly enriched proteins in GBM EVs (Metascape). MCODE 1 in red (R-HSA- 1799339, CORUM:306, WP477), MCODE 2 in blue (Hsa04141, GO:0034976, R-HSA- 901042), , MCODE 3 Green (R-HSA-5663213, R-HSA-3928662, R-HSA-8856828), MCODE 4 in Purple (GO:0034975, GO:0030433, GO:0036503) and MCODE 5 in orange (R-HSA-5627117, R-HSA-5625900, R-HSA-416572).

**Figure 5.**
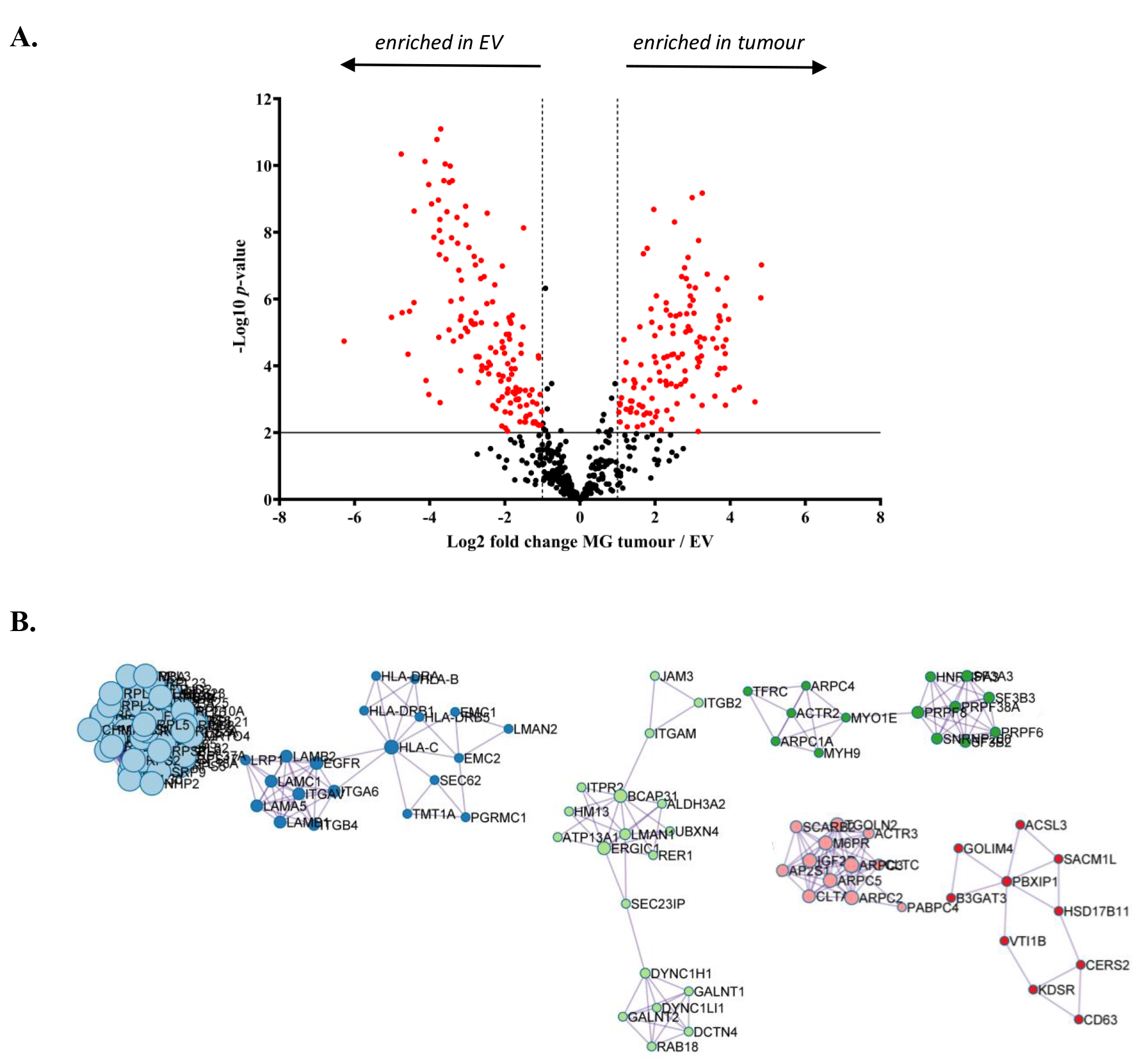
Analysis of proteins common between MNG tumours and their EVs. (A). Volcano Plot illustrating 97 proteins significantly enriched in MNG EVs. Comparison of tumours and the derived EV was performed and tested with Welch’s T-test with post-hoc Benjamini-Hochberg FDR multiple comparison correction (FDR was set at 0.01). Volcano plot was generated using GraphPad Prism 9. (**B).** MCODE analysis of exclusive significantly enriched proteins in MNG EVs (Metascape). MCODE 1 in light blue (CORUM:306, WP477, R-HSA-156902), MCODE 2 in dark blue (M158, R-HSA-3000157, CORUM:2319), MCODE 3 in light green (R-HSA-6811436, R-HSA-8856688, R-HSA- 6811442), MCODE 4 in dark green (CORUM:1181, R-HSA-72163, R-HSA-72172), MCODE 5 in pink (R-HSA-8856828, R-HSA-199991, R-HSA-5653656) and MCODE 6 in red (GO:0008610, R-HSA-556833).

We next determined the differences in the abundance of the 566 proteins shared by MNG tumours and their EVs (**Figure 5A and Supplementary Table 11**). We identified 146 proteins that were significantly upregulated in MNG EVs (**Figure 5A and Supplementary Table 11**). These include HSPG2, NUCB1, MYL6 and PGRMC1 which were found to be associated with EVs isolated from MNG neurospheres (8) (**Supplementary Table 11)**. A protein-protein interaction enrichment analysis was carried out on the 256 exclusive and 146 significantly upregulated proteins in MNG EVs and the subnetworks analysed using the Molecular Complex Detection (MCODE) algorithm in Metascape (33)to identify densely connected network components. Eleven protein complex hubs were identified (**Figure 5B** MCODE 1 - 6). Proteins in MCODE 1 were significantly enriched in ribosome related terms including *ribosome* and *cytoplasmic ribosomal proteins*. Proteins in MCODE 2 were significantly enriched in *lamnin* and *integrin* terms, MCODE3 in *golgi-ER* terms and MCODE 4 in *RNA splicing* terms. MCODE 5 was enriched in *endocytosis, trafficking* and *transport* terms while MCODE 6 was enriched in terns associated with *lipid metabolism*.

### Proteins enriched in EVs relative to tumour type

We next compared the proteome of GBM EVs with MNG EVs to identify exclusive and differently expressed proteins that may be potentially useful for enriching specific tumour derived EV subtypes (**Supplementary Table 12**). A common subset of 591 proteins were present in both GBM and MNG EVs, while there were 291 proteins identified exclusively in GBM EVs and 221 proteins in MNG EVs (**Figure 6A and Supplementary Table 13**). The proteins detected in GBM EVs (and not the MNG EVs) were previously associated with GBM including NEFL, NEFH and MAPT (34) and ATP1A3 and proteins associated with NMDA receptors (GRIA2) (**Supplementary Table 13**). Several cancer stem cell markers were found on GBM EVs including ALDH1, CD44, L1CAM, THY1, CAM2KA and CAMK2B (35, 36) and also CDH2, which was recently identified as a GBM stem cell surface receptor correlating with tumour grade and survival (37, 38). The proteins exclusive to MNG EVs include Talin-1, TPM1, IQGAP1, S100A11, Plastin-2, FBN1, DSP and CALD1, all of which have been associated with MNG previously and shown to be exclusive to MNG relative to GBM (8) (**Supplementary Table 13**).

**Figure 6.**
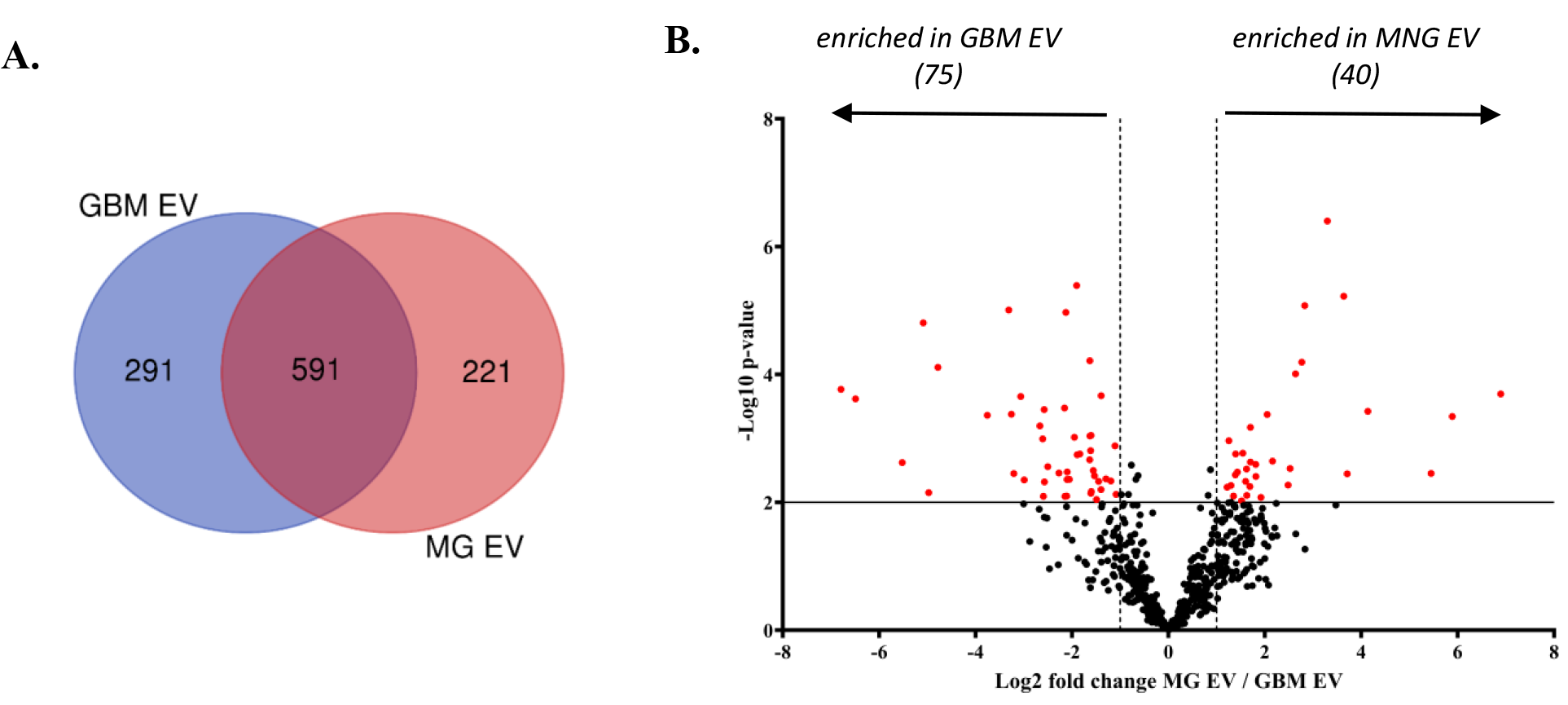
Comparison of GBM and MNG EVs. (A). A two-way venn diagram showing proteins identified in GBM EVs and MG EVs with 291 proteins exclusive to GBM EVs and 221 exclusive to MG EVs. (**B)**. Volcano Plot illustrating 75 proteins significantly enriched in GBM EVs and 40 proteins significantly enriched in MNG EVs. Comparison of GBM and MG EV was performed and tested with Welch’s T-test with post-hoc Benjamini-Hochberg FDR multiple comparison correction (FDR was set at 0.01). Volcano plot was generated using GraphPad Prism 9.

We performed differential expression analyses to determine the relative abundance of proteins common to EVs. We determined the differences in the abundance of the 591 proteins shared by GBM EVs and MNG EVs (**Figure 6B and Supplementary Table 14**). We identified 75 proteins that were significantly up-regulated in GBM EVs and 40 proteins that were significantly upregulated in MNG EVs (**Figure 6B Supplementary Table 15**). Proteins that were up regulated in GBM EVs include GFAP, FSCN1, MYO5A, RCN1 and in MNG include MVP (39), DSP (40), and CKAP4 and AHNAK, which have been associated with particular grades of MNG (41, 42) (**Supplementary Table 15**).

The spatial expression profile of the GBM EV enriched and exclusive proteins (**Supplementary Tables 13 and 15**) examined in an independent RNA-Seq cohort derived from a human GBM tissue cohort (IVY GAP – IVY Glioblastoma Atlas Project) (16). The IVY GAP encompasses a total of 122 RNA samples that were generated from 10 GBM tumours involving a screen of five distinct anatomic regions as identified by haematoxylin and eosin staining. An overview of the normalized gene expression z-score for each corresponding gene from histologically distinct anatomic regions (Leading Edge, Infiltrating Tumour, Cellular Tumour, Perinecrotic Zone, Pseudopalisading Cells and Microvascular Proliferation) is displayed in a heatmap (**Supplementary** Figure 2). Analysis of the positive z-scores for all genes corresponding to the GBM EV enriched and exclusive proteins revealed that whilst that the genes were expressed in all anatomic regions, they were generally highly expressed primarily in the ‘Leading Edge’, followed by the ‘Infiltrating Tumour’ region suggesting GBM EVs contain proteins associated with these pathways regardless of where the location of the GBM.

## Discussion

In this study, employing a method that we developed for isolating EVs from human brain tissue (13, 15, 43), we successfully isolated EVs from frozen human tumour tissue biopsies obtained during surgery from patients diagnosed with either GBM or MNG. This is the first report of in situ EVs isolated from brain tumours. We determined the protein content of the tumours and their EVs using data dependent analysis tandem mass spectrometric proteomics. This allowed us to identify proteins that were exclusively detected or enriched in EVs relative to the corresponding tumour samples.

Extracellular vesicles derived from both healthy and tumour cells encompass a wide array of molecular components including nucleic acids, fragments of organelles, lipids, metabolites and protein. The precise mechanisms governing the selection of molecules for inclusion in EV cargo are still under investigation, however proteins found within EV have been reported to be associated with progression of various cancers including GBM (1, 2). The detection of GBM EV exclusive proteins (not detected in the corresponding tumour tissue), included EV biogenesis proteins such as CD63, SDCBP, VAMP2 and CD44A. Differential expression analysis was utilized to reveal the relative abundance of proteins that were enriched in the EVs compared to the tumour tissue. Our study not only confirms the presence of proteins previously identified in GBM EVs from various sources (including CUSA washings, cell lines, and plasma)(4, 7–12, 25), but also crucially identified novel proteins that have not been reported before in association with GBM EVs before including proteins associated with solute carrier transporters and fatty acid transport. Recently it has been shown that solute carrier family members can promote the progression of GBM by activating the PI3-AKT signalling pathway (44). Proteins which function as fatty acid transporters can be used to generate ATP via fatty acid oxidation (FAO) (45), which in turn can contribute to GBM invasion (ref) and proliferation (46)during tumour progression These proteins could therefore potentially be EV markers of tumour progression or recurrence after surgery.

Meningiomas are the most common primary brain tumour, constituting 38% of all central nervous system tumours (47). The primary treatment for these tumours is surgical removal. Around 20% of these tumours are prone to aggressive behaviour (which is not always accurately predicted by the WHO classification), highlighting the need for more refined prognostication methods. These aggressive tumours contribute significantly to patient morbidity and mortality (48, 49). Moreover, the development of a readily available and dependable biomarker would greatly benefit ongoing patient management. Current follow-up procedures heavily depend on regular MRI scans, which are not only costly and labour-intensive, but also fail to offer insights into the molecular changes that signal a transition to a more aggressive tumour phenotype. Analysing MNG EV profiling (tumour or blood) could help identify aggressive tumours and guide post-operative adjuvant therapy.

While there have been several studies on GBM derived EVs, only a few have been conducted on MNG derived EVs (7–9, 18, 50, 51). A study by Ricklefs et al (2022) observed that the levels of circulating EVs in the blood of meningioma patients are increased, correlating with malignancy grade and oedema, but subsequently declined post-surgery (8). Proteomic profiling of the EVs (patient plasma or MNG cell lines) identified 77 proteins exclusive to the meningioma EVs, including the meningioma marker, desmoplakin (52). We compared our exclusive meningioma EV proteins (221) with the 77 proteins in the Ricklef study and identified 12 common proteins which could be potentially useful for enriching meningioma EVs - aminopeptidase N, caldesmon, C-type mannose receptor 2, desmoplakin, EGF-containing fibulin-like extracellular matrix protein 1, fibrillin-1, plastin-2, Ras GTPase-activating-like protein IQGAP1, talin-1, tenascin-X, transforming growth factor-beta-induced protein ig-h3 and transgelin.

Hader et al (2024) performed a large-scale targeted proteomics study of 51 serum and 53 fresh frozen tumour samples from low- and high-grade meningioma biopsies (53). A total of 15 proteins from tissue and 12 proteins in serum were found to be the best segregators using a feature selection-based machine learning strategy with an accuracy of around 80% in predicting between low grade (grade I) and higher grade (grade II and II) meningiomas. When these were compared against the MNG EV proteins we identified, a total of 11 common proteins were identified (PGK1, ALDOA, ANAXA1, HSPB1, PFN1, PKM, S100A11, SPTBN1, SPTAN1, PLEC and TF). Our study not only confirms the presence of proteins previously identified in MNG EVs(4, 7–12, 25), but also identified novel proteins that have not been previously reported in association with MNG EVs.

In comparing GBM EVs with MNG EVs, we identified the 291 exclusive and 75 enriched proteins in GBM relative to MNG EVs. Assessment of the IVY GAP gene expression revealed that these proteins are more highly expressed at the leading edge of a GBM, where tumour cells can form invadopodia which degrade the ECM facilitating invasion into the surrounding healthy brain parenchyma. Importantly, analysis of our exclusive and enriched proteins in GBM EVs in the Glioma Bio Discovery Portal (38) demonstrated that a higher expression of these proteins had a significant impact in reducing GBM patient survival. We have previously demonstrated that GBM cells can form invadopodia to facilitate GBM cell invasion (54) and that extracellular vesicles can promote invadopodia activity in recipient GBM cells *in vitro*, in response to their increased secretion post-radiation and temozolomide treatment (11).

Our study demonstrates that *in situ* EVs can be isolated from the ECM of human brain tumours. Analysing the cargo of these EVs can provide insights into their possible function. We discovered a set of proteins that are exclusively enriched in GBM EVs that could possess prognostic potential, are potentially more highly expressed at the leading edge of tumours and are linked to invadopodia, which would be present at the tumour edge facilitating tumour cell invasion into the surrounding brain. Future studies using larger sample sizes and analysing different tumour types and grades is warranted.

## Acknowledgements

The authors acknowledge the use of the transmission electron microscope at the Ian Holmes Imaging Centre, The University of Melbourne.

---

**Supplemental Figure 1.**
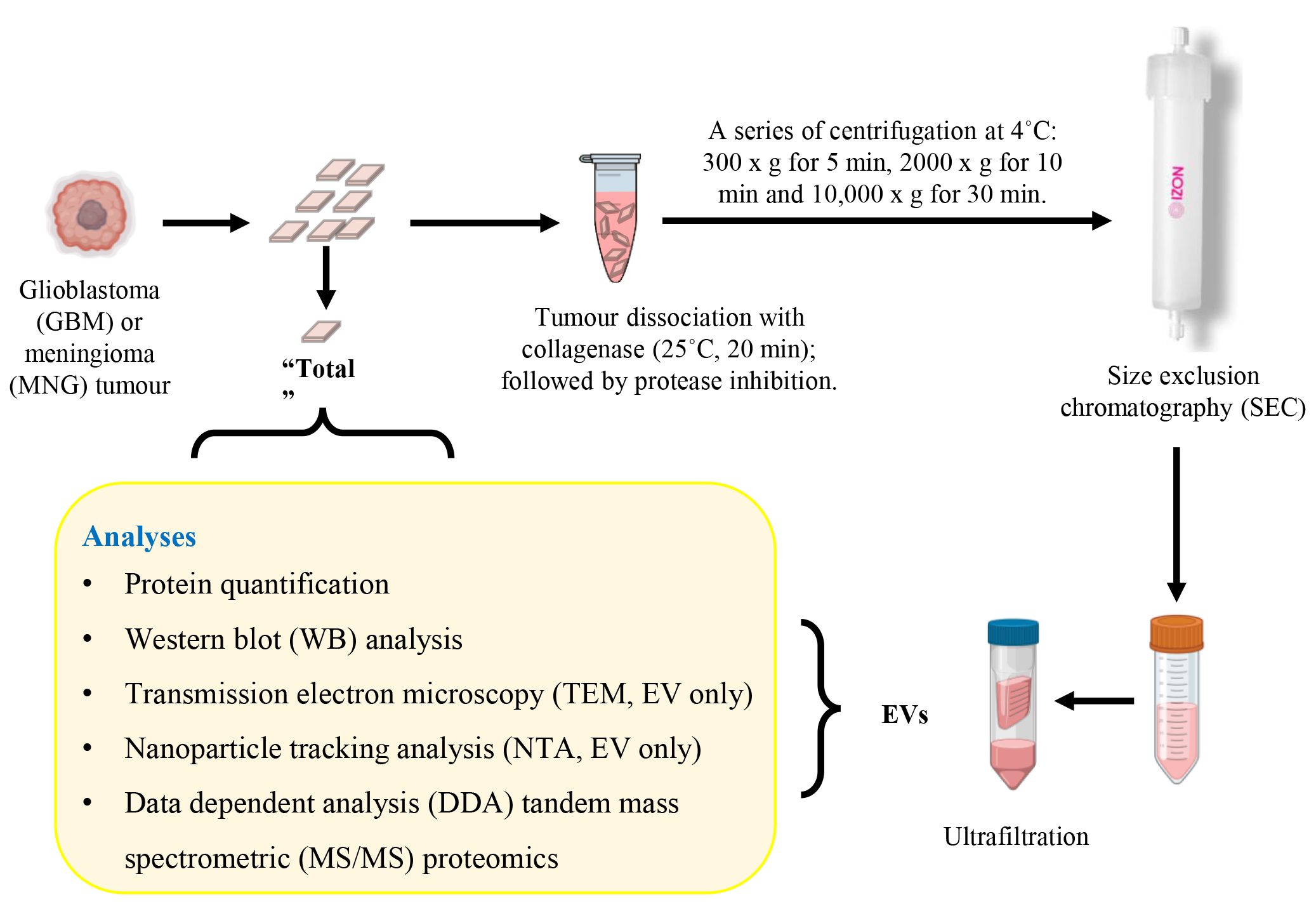
Summary of the experimental workflow for human post-mortem tumour EV isolation and analysis. Fresh frozen glioblastoma or meningioma tumour tissue (n=8 each) were sliced on ice. A small section of each was collected as a sample control (“Total”). The remaining sections were dissociated with 50 U/mL of collagenase type 3 in DPBS at 25°C for a total of 20 min with shaking, followed by addition of protease and phosphatase inhibitors and EDTA. The dissociated tissue was spun at 300 x g for 5 min at 4°C and the supernatant was spun at 2,000 x g for 5 min at 4°C, followed by a 10,000 x g spin for 30 min at 4°C. The extracellular vesicle (EV) containing 10,000 x g supernatant was loaded on to the qEV10 70 nm SEC column and the EV elution (20 mL) after the first 20 mL void volume was collected and concentrated using a 10kDa centrifugal filter. The tumours and EV were subjected to biochemical and biophysical analyses.

**Supplementary Figure 2.**
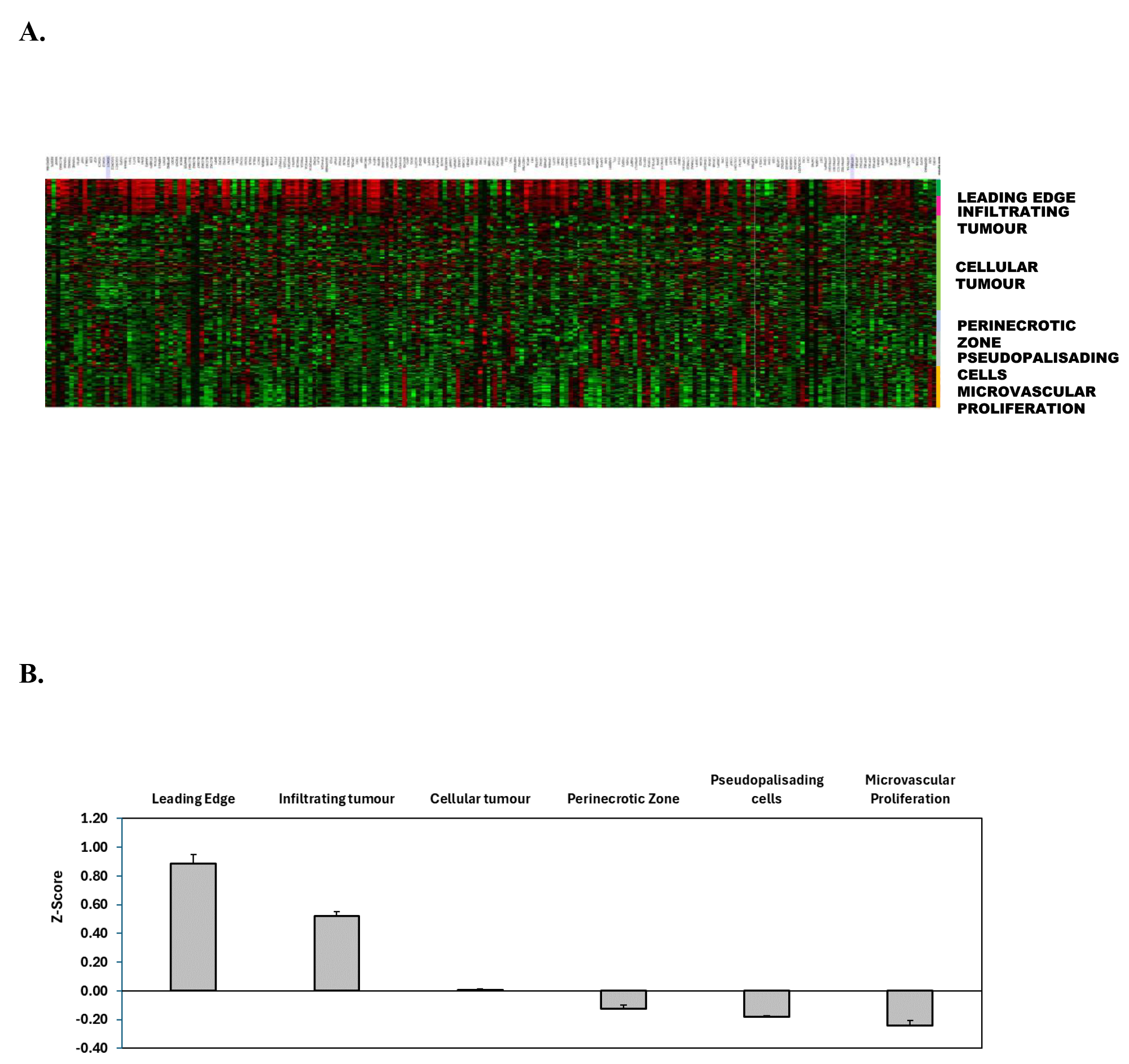
**A.** Normalized RNA sequencing data from the IVY GAP database showing expression of the corresponding genes to GBM exclusive and enriched proteins (relative to MNG EVs) in the histologically distinct anatomic regions (Leading Edge; Infiltrating Tumor, Cellular Tumor, Perinecrotic Zone, Pseudopalisading cells and Microvascular Proliferation). The ‘Leading Edge’ and ‘Infiltrating Tumor’ regions show that many samples display a positive gene expression ‘z-score’ (red colour). **B .** Graphical presentation of the average z-score for each anatomical region (utilizing the z-score for each gene within the anatomical region).

## Supplementary Material

*Supplementary* Figure 3. **(A)** Expression heatmap of the corresponding genes to GBM exclusive and enriched proteins (relative to MNG EVs) using data from the 3-platform analysis in the TCGA subtype classification of 200 GBM patients . Colour annotations are according to profile similarity, subtype classification and prognostic index stratification. A list of 50 or more molecules are filtered to keep up to the 50 most varied molecules amongst the samples. Molecule names are annotated with Hazard Ratios (HR) from Cox analysis; * indicates HR’s with p-vals < 0.1; ** indicates HRs with p-vals <0.05. (**B)** Survival analysis based upon the impact of the multi-gene prognostic index. A Cox proportional-hazards regression model analysis of the corresponding genes to the GBM exclusive and enriched proteins from above, as a covariate and stratification down the median was performed for the Full cohort of GBM patients, as well as according to their GBM subtype. GBM patients with a Classical (C-subclass), Mesenchymal (M-subclass), or Proneural (P-subclass) profile and lower expression of the genes are associated with a significant improvement in survival. The Cox-model for GBM patients with a Neural (N-subclass) could not be computed, as it is non-convergent (NC). Importantly, a lower expression of the genes across the Full cohort of GBM patients is also associated with a significant improvement in survival.

*Supplemental Tables 1 - 15.* Subject case information and proteomic data.

